# Food-Derived Compounds Extend the Shelf-Life of Frozen Human Milk

**DOI:** 10.1101/2024.12.11.627965

**Authors:** Justin E. Silpe, Karla Damian-Medina, Bonnie L. Bassler

**Affiliations:** Department of Molecular Biology, Princeton University, Princeton, New Jersey, USA; Howard Hughes Medical Institute, Chevy Chase, Maryland, USA; PumpKin Baby Inc., Princeton, New Jersey, USA

## Abstract

Breastmilk is known to provide optimal nutrition for infant growth and development. A cross-sectional analysis of nationally representative US data from 2016 to 2021 revealed that >90% of lactating mothers reported using breast pumps to express milk.^1^ We conducted a survey of *n* = 1,049 lactating or recently lactating individuals from a US nationally representative population to explore breastmilk storage practices among this group. The data revealed that 83% of respondents store breastmilk in their homes, with 68% using freezers to do so for >1 month. The lowest available temperature in most household freezers is -20 °C, a temperature that is inadequate to maintain human milk’s emulsified structure, leading to separation, degradation of fats, loss of key vitamins, and changes in palatability. We developed a first-of-its-kind high-throughput screening platform to identify food-derived compounds and combinations of compounds that, when added to human breastmilk, preserve fat content, retain antioxidant capacity, and reduce production of rancid-associated free fatty acids during extended freezer storage. These formulations represent leads for the development of safe and affordable frozen breastmilk shelf-life extenders for easy at-home use to increase the longevity of stored breastmilk.

## Introduction

Human breastmilk is the gold standard for infant nutrition, containing a complex blend of vital nutrients and other factors that cannot be synthesized or sourced from other mammals, yet are essential to meet life’s critical early milestones. The World Health Organization (WHO) recommends infants be exclusively breastfed for the first 6 months of life. However, fewer than half of all infants currently meet this guideline,^2^ and the majority of parents (60%) fail to accomplish their own breastfeeding goals.^3^ In today’s work-life system, many new parents are full-time employed at the time of childbirth, and most return to their professions while their infants still require breastmilk.^4^ Alarmingly, in the US, 43% of women leave the workforce within 3 months of childbirth,^5^ and those who remain earn ∼30% less in subsequent years than they did the year before the birth of their first child.^6^ One reason cited for leaving the workforce is the desire to take care of the infant, for which feeding the baby is one of the highest-ranking needs.^7^ This desire often conflicts with the demands of returning to work, as evidenced by the fact that among the 67% of working mothers who initiated breastfeeding in one study, only 10% continued once back to work.^8^

Given the critical role breastmilk plays in infant development, many parents rely on freezer storage to enable continued breastfeeding. However, household freezers fail to prevent separation of milk’s emulsified components, and freezing can accelerate the degradation of proteins, vitamins, and other bioactive compounds.^9–14^ The structural changes induced by freezing can alter milk palatability, potentially leading to infant rejection.^15^ In fresh human milk, lipids are encapsulated within milk fat globules (MFGs), which are surrounded by fat globule membranes, termed milk fat globule membranes (MFGMs). MFGMs protect lipids from lipases present at the globule interface and/or in the aqueous phase. The crystallization of milk lipids that occurs during freezing can damage MFGMs, allowing lipases access to lipids. Lipases hydrolyze milk triglycerides into free fatty acids (FFAs) and glycerol. FFAs, particularly volatile short and intermediate-chain fatty acids, coincide with rancid flavors in milk products.^15^ Indeed, FFA production affects flavor and accelerates oxidative processes that further degrade sensory and nutritional milk quality. These issues are magnified by the fact that most home cooling systems are subject to temperature fluctuations and variable personal habits,^16^ further degrading milk quality. Consequently, parents often face the dilemma of discarding stored milk or feeding their infants breastmilk they suspect is compromised.

Evidence supporting the changes human milk undergoes following freezing includes a study reporting that 25% of infants refused to consume previously-frozen human milk, and in 95% of these cases, the milk was described as smelling “off”.^17^ In the current work, we refer to off-smelling milk as being rancid. Rancidification is a process in which fats undergo oxidation, autoxidation, hydrolysis, and/or lipolysis upon exposure to air, light, moisture, or enzymes.^18–21^ Rancid milk, while still nutritious and safe to consume,^22^ is characterized by an unpleasant taste and odor. Many parents are loath to feed rancid milk to their infants and not all babies accept such milk, suggesting that these changes prevent some infants from receiving any nutritional benefit from breastmilk. Discarding or rejection of breastmilk forces suboptimal feeding regimes for parents (i.e., resorting to commercial formulas despite knowing them to be imperfect substitutes for breastmilk). Compounding this problem is that current methods to facilitate milk storage such as freeze-drying, pasteurization, and scalding have limitations, including high costs, loss of nutritional content, and impracticality for home use. Therefore, innovative approaches that can mitigate the adverse effects freezing has on breastmilk, while remaining affordable and accessible, are essential. In this work, we sought to understand current breastmilk storage practices and challenges encountered. Based on the data acquired, we propose safe and easy to use solutions.

## Results

### Breastmilk Freezer Storage Respondent Demographics and Practices

It is generally understood, particularly in developed countries with limited parental leave policies, that nursing parents depend heavily on storage of expressed breastmilk. However, challenges associated with pumping and the consequences of at-home storage on the nutritional quality of human milk have not been thoroughly quantified. To address this gap, we conducted an anonymous, electronic, retrospective survey to garner data on current breastmilk storage practices, the hurdles faced, and the consequences of the use of pumped human milk as reported by parents. The survey was conducted on a nationally representative sample of 1,049 reproductive-age individuals across 50 states who had considered, attempted, or successfully breastfed an infant within the past 3 years (see survey criteria in Supporting Information and SI Fig 1a-f). The survey asked participants to indicate the methods used to feed their infants during the first 6 months of life. Forty-six percent reported feeding their infant exclusively breastmilk, while 11% used only formula, and 44% fed their infants a combination of breastmilk and formula (SI Fig 1a). Regarding how breastmilk was provided to infants, 55% of respondents reported using a combination of both breastfeeding and pumped milk, 26% reported that breastmilk was delivered exclusively from the breast, and 19% exclusively pumped and bottle-fed breastmilk (SI Fig 1b). The average age of participants who completed the survey was 30.6 ± 6.7 years old. Sixty-one percent were White, 23% were Black or African American, 17% were Hispanic or Latino, 4% were Asian, 2% were Native American or Alaska Native, and 1% were Middle Eastern or North African. Forty-four percent of participants were full-time employed, 19% were part-time employed, 22% were unemployed, and 4% were full- or part-time students. Approximately 81% of participants reported having an annual household income between USD 20,000 to 199,000/year (Supplemental Table 1).

**Figure 1.**
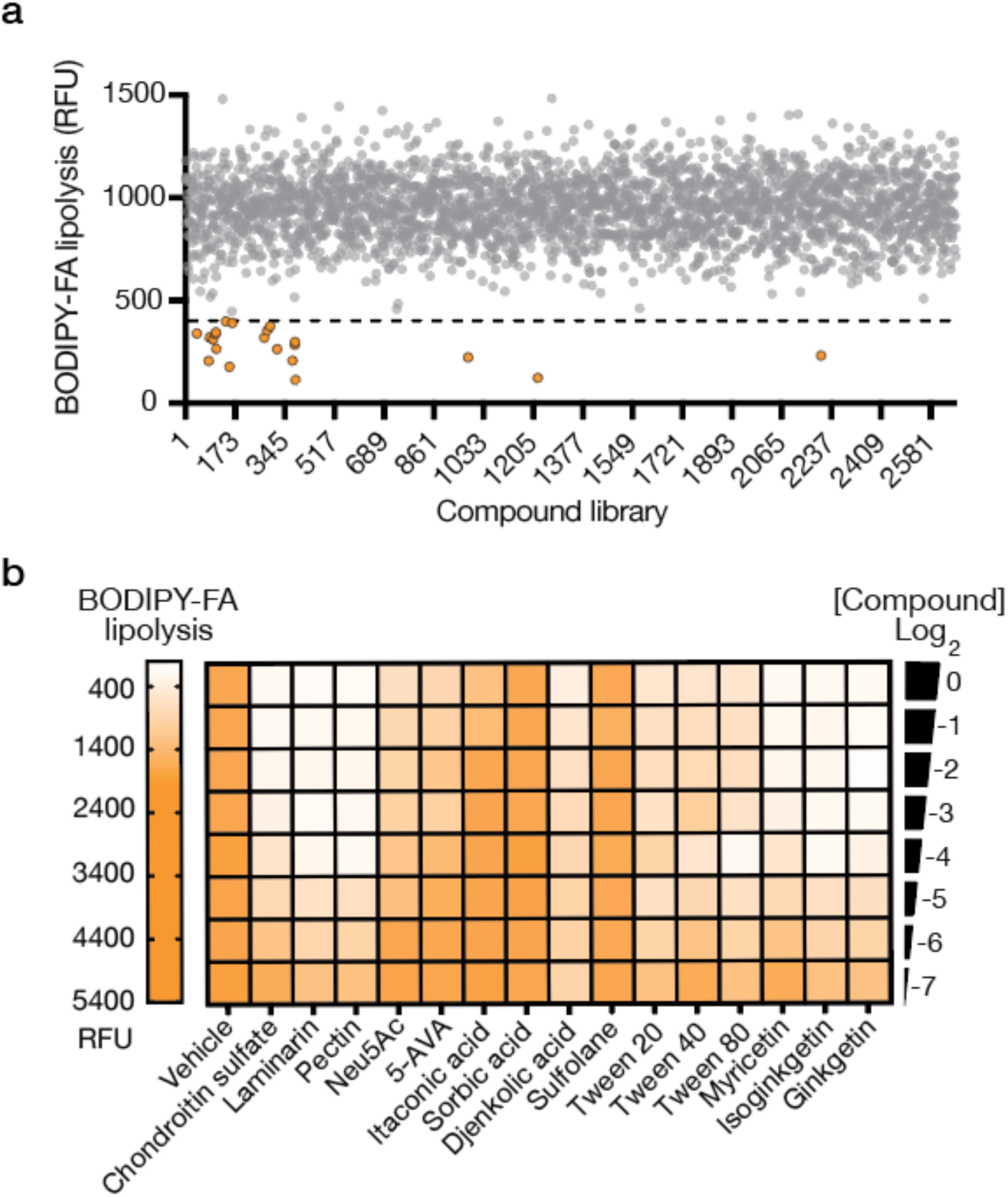
High throughput screening and validation of compounds that stabilize breastmilk by reducing lipolysis. **(a)** Food-derived compounds screened for breastmilk lipolysis inhibition. Orange dots indicate compounds reducing lipolysis below the threshold (dashed line); gray dots show inactive compounds. **(b)** Validation of selected hit compounds from panel a. The heatmap displays dose-dependent relative fluorescent units (RFU). Compound concentrations are expressed as dilutions (Log2(0) to Log2(-7)) of the screening library stock solutions. Ne5Ac: N-Acetylneuraminic acid; 5-AVA: 5-Aminovaleric acid. Data represent the initial HTS readout (a) and the median of three independent replicates (b).

The survey responses confirmed that storage is a common practice among parents who use breastmilk. Specifically, 83% of respondents reported having stored breastmilk, of whom 68% reported using a freezer to do so (Supplemental Table 2). The duration of storage in freezers varied, with 31% of respondents storing milk for less than 1 month, 40% for 1 to 3 months, 17% for 4 to 6 months, and 12% for more than 6 months (SI Fig 1c). Reasons for freezing breastmilk were related to oversupply (63%), work (38%), and travel (23%) (Supplemental Table 2). The survey also revealed that surprising amounts of breastmilk go to waste. Specifically, 75% of respondents reported discarding breastmilk, primarily due to incomplete feeding (62%), questionable proper storage (38%), and/or sensory changes (i.e. “the milk smelled off,” 23%) (SI Fig 1d). Changes in smell or taste following defrosting were reported by nearly 20% of respondents (Supplemental Table 2). These sensory alterations presumably affect the acceptability of stored milk by infants, as 26% of respondents reported experiencing occasional or persistent infant rejection of the milk (SI Fig 1e). Moreover, the proportion of respondents reported that changes in the milk increased in a storage-time dependent manner (SI Fig 1f), suggesting that at least some of the observed alterations are driven by the storage process itself rather than by interindividual differences in breastmilk composition or variability in human practices.

### Changes that Occur in Human Milk During Storage

With the goal of improving the shelf-life of stored human milk, we devised a high-throughput, small-molecule screening (HTS) approach to identify compounds naturally present in foods that stall the rancidification process. To verify that lipolysis and lipolytic byproducts can be reliably measured in small volumes of human milk, we adapted existing assays for use with human milk. One assay employed a fluorescent fatty acid conjugate consisting of a 4,4 difluoro-methyl-4-bora-3a,4a-diaza-s-indacene-3-dodecanoic acid (BODIPY) dye and a 4,4 dimethylamino-phenyl-azo-benzoic acid (DABCYL acid) quencher to quantify active lipolysis at the point of testing (BODIPY-FA). Second, a coupled-enzyme assay was used to measure glycerol, a product of triglyceride metabolism, as a proxy for fat breakdown. This assay couples glycerol metabolism to NADH production, which drives a luciferase reaction generating bioluminescence emission proportional to glycerol concentration (Glycerol-glo). SI Fig 1g (upper), shows that the rate of lipolysis generally decreased during human milk freezer storage but remained measurable. Moreover, the concentration of glycerol increased across all freezer storage time points (SI Fig 1g, lower). These data demonstrate that lipolysis and the accumulation of lipolytic byproducts can be measured as they occur in breastmilk. Our assays are quantitative, scalable (96 and 384-well plate formats), and adapted for low milk volumes (2 μL per condition tested), making the strategy amenable to HTS. HTS is a cornerstone approach used in drug discovery and has broad applications in chemistry and biology, but to our knowledge, it has never been applied to breastmilk.

### High-Throughput Screening of Food-Derived Compounds for Extending Stored Human Milk Shelf-Life

We screened approximately 2,750 food-derived compounds from three libraries (Biolog Phenotype Microarrays, MCE Food Additive Library, and MCE Food-Source Compound Library) for the ability to attenuate lipolytic activity, as measured by BODIPY-FA, after 1 week of household -20 °C freezer storage. Our screening process produced 21 hits, defined as yielding

<400 relative fluorescent units from the BODIPY-FA probe (Figure 1a), 15 of which repeated upon retest. We assessed the 15 candidate hits for their abilities to reduce lipolysis and FFA accumulation after 3 months of freezer storage (Figure 1b and SI Fig 1h, left, respectively). Importantly, we demanded that the compounds did not suppress lysozyme and protease activity, two important, non-fat related enzyme activities known to be present in human milk (SI Fig 1h, middle and right, respectively). We defined lysozyme and protease retention as >75% of the levels in fresh or untreated breastmilk. Using these criteria, we identified nonionic surfactants, glycosaminoglycans, pectins, and specific biflavonoids as hits of interest. Based on these data and, as proposed in our model (see Discussion), we hypothesize that the compounds slow the degradation of breastmilk by stabilizing its structural integrity without inhibiting important enzymes that the infant requires upon ingestion.

In human milk, endogenous ascorbic acid (i.e., vitamin C) levels reportedly decrease by 20% after 24 hours of refrigeration,^23^ with ascorbic acid declining to undetectable levels in some samples after 2 months of freezer storage.^24,25^ Supplementation of human milk with exogenous ascorbic acid (100 μg/mL) preserves antioxidant capacity following 1 month of household -20 °C freezer storage (Fig 2a). To test whether the same effect occurs in the presence of lipolysis-suppressing food-derived compounds, we supplemented human milk with ascorbic acid and hit compounds from the screen. Fig 2a shows that, after 1 month of household -20 °C freezer storage, we achieve additive effects on preservation across combinations. Specifically, ascorbic acid (100 μg/mL) addition did not slow lipolysis, yet it maintained antioxidant capacity. Conversely, a compound identified from our lipolysis-inhibiting screen, pectin (0.5% w/v), had a limited effect on antioxidant maintenance but effectively curbed lipolysis (Fig 2a). We prioritized pectin for further investigation given its extensive clinical history as a safe dietary product for toddlers and children.^26–28^ Our findings suggest that combining compounds that influence distinct properties of the milk can maintain multiple qualities related to breastmilk nutritional value, and thereby significantly improve breastmilk preservation.

**Figure 2.**
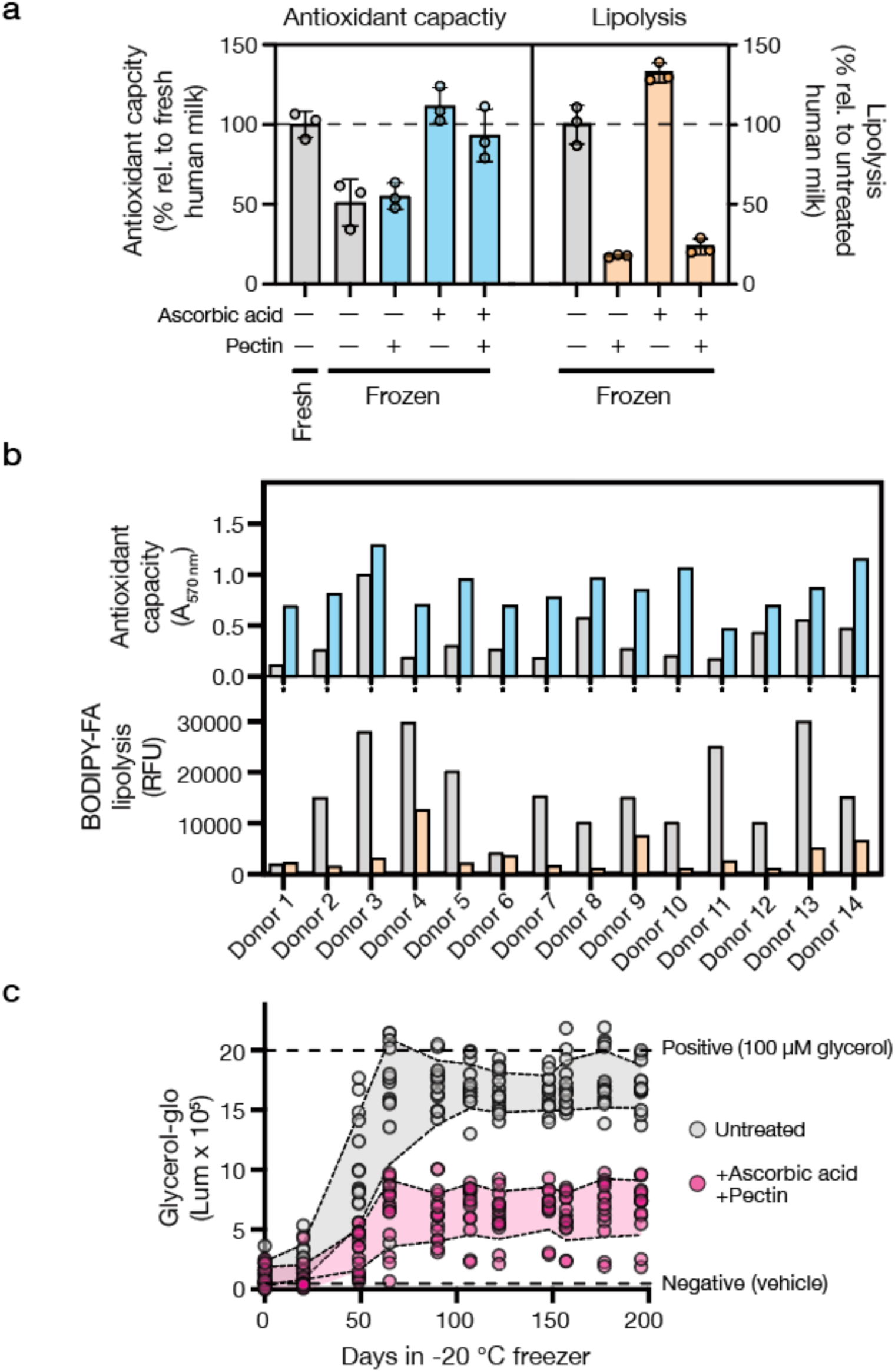
Effects of ascorbic acid and pectin on antioxidant capacity, lipolysis, and glycerol levels in frozen human milk. **(a)** Antioxidant capacity and lipolysis in milk treated with 100 μg/mL ascorbic acid and/or 0.5% w/v pectin. No treatment (–) or the designated treatment (+) is indicated under the x-axis. Values are normalized to untreated fresh milk (antioxidant capacity) or untreated frozen milk (lipolysis). Gray bars: untreated samples; colored bars: treated samples (blue: antioxidant capacity; orange: lipolysis). **(b)** Donor variation (*n* =14) in antioxidant capacity (A_570_ nm, blue) and lipolysis (BODIPY-FA RFU, orange) with and without the combined 100 μg/mL ascorbic acid and 0.5% w/v pectin treatment. Gray: untreated; colored: treated samples. **(c)** Glycerol accumulation during -20 °C storage of human milk with and without the combined 100 μg/mL ascorbic acid and 0.5% w/v pectin treatment. Dashed lines indicate positive (100 μM glycerol) and negative (vehicle) controls. Data show mean ± std of three independent replicates (a), individual donor measurements (b), and mean ± std of 14 biological replicates (c).

The exact chemical composition of expressed human milk is known to be person-specific and, within a single individual, varies over time, making it difficult to predict if or when stored expressed human milk will become rancid. To test the generalizability of our approach, we expanded our sample size to 14 donors who provided freshly expressed milk. Fig 2b demonstrates that despite wide inter-individual variability in baseline lipolysis activity, addition of 100 μg/mL ascorbic acid and 0.5% w/v pectin prior to freezing proved effective in reducing fat breakdown and increasing antioxidant capacity across donors following 3 months of freezer storage. Moreover, across donors, the combination of ascorbic acid and pectin eliminated, on average, approximately 60% of the glycerol production that normally occurs in frozen breastmilk after more than 6 months of freezer storage (Fig 2c).

## Discussion

### Implications of Freezer Storage on Breastmilk Quality, Innovative Approaches to Breastmilk Preservation, and Study Limitations

Most household freezers cool to -20 °C, a temperature that does not adequately preserve the complex emulsified structure of human milk. Moreover, while -20 °C is generally the lowest setpoint for home freezers, a recent report on at-home refrigerated insulin indicates that, in practice, domestic cold-storage conditions rarely meet recommended temperatures (78.8% of refrigerated insulin fell outside of range at least once, with the average insulin being out of range for 3 hours per day).^16^ Consistent with this finding, it has been reported that storing human milk under suboptimal freezer conditions can lead to significant alterations in its nutritional composition. Specifically, essential macronutrients such as proteins, carbohydrates, and lipids undergo accelerated degradation. Lipids are particularly affected, displaying a reduction of approximately 3.9% after 7 days and 9% after 3 months of storage.^12,29^ Key micronutrients, including vitamins, are also negatively affected following freezing. Lastly, prolonged freezer storage causes milk’s endogenous bactericidal properties to decline.^30–36^

Germane to the current report, human milk contains antioxidants, including enzymes, vitamins, and non-enzymatic proteins, whose concentrations are affected by storage conditions.^37–39^ Studies on human milk have shown that antioxidant capacity decreases during both refrigeration and freezing, with freezing causing a larger reduction.^40^ Losses in antioxidant capacity are directly related to rancidification: FFAs released from lipolysis are highly susceptible to oxidation, leading to the formation of lipid peroxides and secondary oxidation products, such as aldehydes and ketones that impart off-flavors and odors.^41,42^ The slow freezing rate and variable temperatures of household freezers, and particularly the freeze-thaw process, are known to be damaging, especially to emulsified foods, because of differences in the freezing and melting points of their fat and water phases. Consequently, in the context of breastmilk, the freeze-thaw process drives phase separation which destroys milk’s natural emulsified structure.^43,44^ Phase separation promotes non-specific enzymatic activities, changes in substrate availabilities, and production of reactive intermediates at concentrations that vastly differ or do not occur under physiologically relevant conditions (37 °C).^45^ We propose that losses in structural integrity and changes in constituent composition combine to drive breastmilk rancidification (Figure 3a).

**Figure 3.**
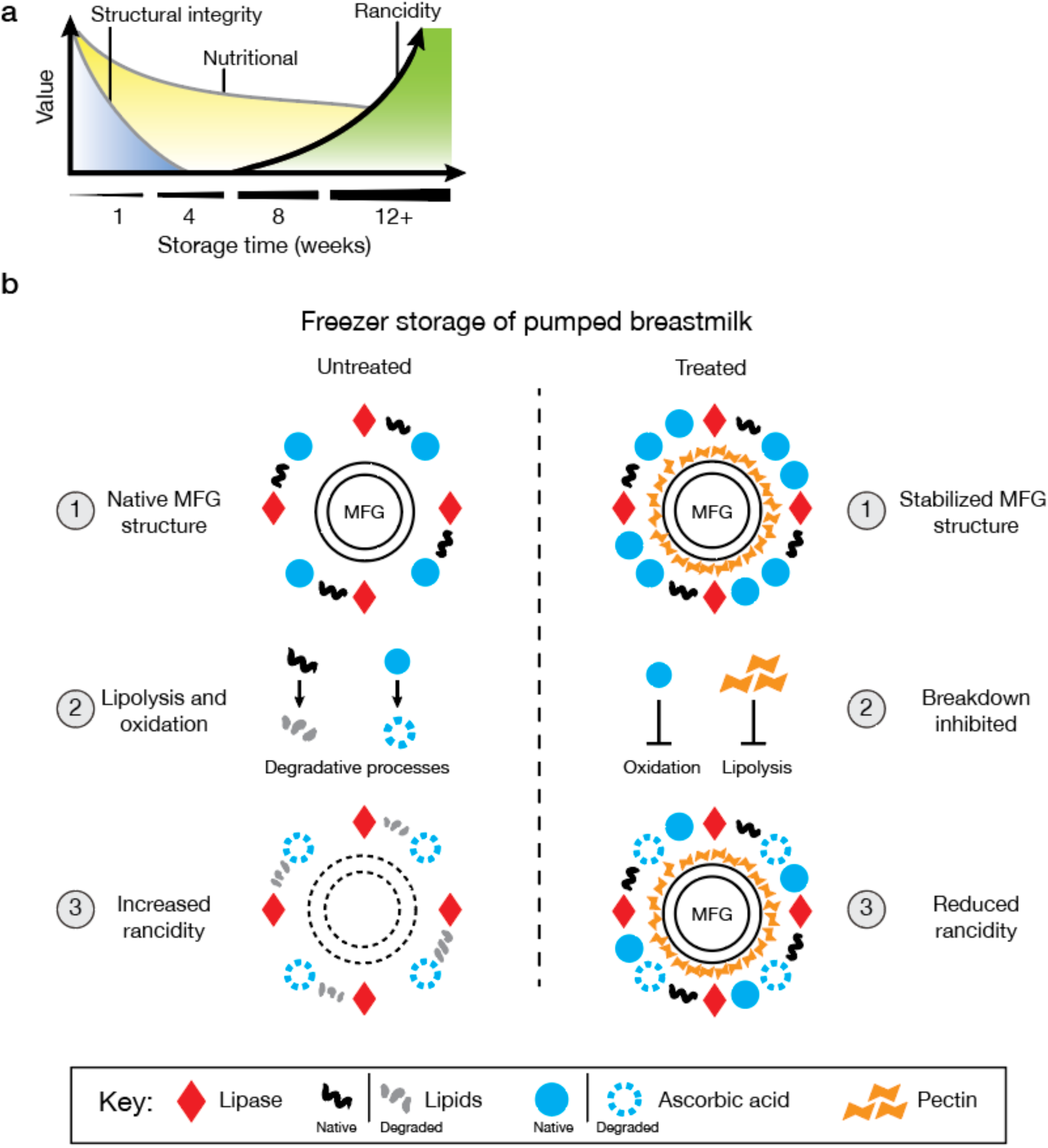
Changes in breastmilk quality during storage and the protective effects of ascorbic acid and pectin treatment. **(a)** Model depicting changes in human milk quality over storage time. At 1 week, degradation begins, leading to a decline in both nutritional value and structural integrity. Between 8 to 12 weeks, rancidification accelerates, marked by an increase in FFAs and other breakdown products, which further reduce nutritional quality and structural integrity. Rancidity continues to rise beyond 12 weeks. **(b)** Schematic model illustrating biochemical and structural changes that occur in human milk during storage. Untreated milk, left: Storage compromises MFGM structure, enabling lipase-mediated fat hydrolysis and degradation. Lipolysis of triglycerides and lipid oxidation produces secondary and tertiary products that cause rancidity. Treated milk, right (model proposed in the current work): Pectin preserves MFGM structural integrity, limiting lipase access to MFGMs. Ascorbic acid provides protection from oxidation. Together, these two mechanisms suppress rancidification.

To address the issue of rancidification, we suggest exploiting food-derived compounds to stabilize the structural integrity of breastmilk during freezer storage (Figure 3b). Our hypothesis centers on the protective roles of ascorbic acid and pectin, compounds identified in this work, to preserve the structure and nutritional quality of human milk by stabilizing the emulsion and, in so doing, inhibiting oxidation. Fats in human milk are contained within MFGs, delicate fluid structures enveloped by membranes derived from mammary epithelial cells from the mother’s mammary gland (i.e., MFGMs). As reaction sites, MFGs and MFGMs carry extremely high local concentrations of fats. During storage, particularly under freezing conditions, the structure of the MFGM is compromised.^46^ As shown in Figure 3b, freezer storage enables lipases (red diamonds) to access and hydrolyze fats, releasing FFAs (black squiggles). Oxidation of unsaturated fatty acids leads to the formation of primary products, including peroxides, which degrade into secondary products such as aldehydes and unsaturated alkenals. These compounds further react to form tertiary degradation products, including aldol condensation products, hydroxy alkenals, and alkyl furans, contributing to rancidity.^47^ We propose that pectin interacts with MFGMs, as has been shown for lipid droplets,^48^ by forming a surrounding stabilizing layer via surface activity or electrostatic interactions with ionized carboxylic groups that prevent MFG coalescence and enhance emulsion stability (Figure 3b). Regarding oxidation, considering both safety and historical use in infants, ascorbic acid is an ideal antioxidant that is naturally present in all freshly expressed human milk and is an ingredient in commercial infant formulas. While ascorbic acid is a commonly used food additive, it is known to degrade rapidly in a storage-dependent manner.^49^ Indeed, to account for expected losses, commercial infant formula- and first food-makers routinely include significant overages of vitamin C. For example, a formula labeled with 8 mg/100 kcal vitamin C may actually contain >400% of that amount, depending on when it is measured.^50^ In our system, the addition of ascorbic acid presumably extends breastmilk shelf-life by increasing the amount and longevity of antioxidant capacity, which combats FFA oxidation.

A significant attribute of our method is its reliance on naturally occurring compounds rather than potent small-molecule inhibitors that disrupt enzyme activity, such as orlistat, a broad-spectrum lipase inhibitor.^51^ By avoiding such inhibitors, we aim to preserve the milk’s natural enzymatic processes and ensure that the nutritional integrity and digestibility of breastmilk remain intact.

We note that the compounds we have identified have not been evaluated by a regulatory body for use in this application, underscoring the need for further research and development. Additionally, our study has several experimental limitations including, in the national survey data, the possibility of self-reporting biases; and, in the milk collection effort, the donor pool was confined to New Jersey for the practicality of collecting fresh samples. Despite these limitations, our findings demonstrate the potential of our formulation to preserve nutritional value and reduce fat breakdown in freezer-stored breastmilk. We are now evaluating the benefits of our formulation on other high-value biological components present in frozen-stored human milk and, if warranted, we will refine the ingredients to support preservation of additional milk components. Future research will aim to validate our findings through larger, more diverse cohorts and longer-term studies to verify the efficacy and safety of the proposed preservative formulations.

### The Role of Breastfeeding in Health and Economic Outcomes

Breastfeeding greatly influences health and prosperity yet remains understudied and undervalued, which negatively affects parents’ abilities – particularly women and mothers – to equitably participate in the workforce and the greater economy. At present, infant formula is included in the WHO calculations of global GDP, but breastfeeding is not, which devalues the contributions it has on society.^52^ The WHO has begun the process of quantifying the “work” of breastfeeding for global GDP.^53^ Exclusive breastfeeding is estimated to take 1,800 hours per year of a lactating person’s time, compared to a “full time job” which is estimated at 1,960 hours per year.^54^ Given the time invested, it is alarming that the majority of lactating parents (75% in our survey) discard expressed breastmilk and/or face unsolved hurdles to feeding it to infants after storage (26% in our survey).

While improvements in workplace protection policies and advanced pump technologies continue to reduce barriers associated with the expression of human milk, our work highlights that the nutritional quality of expressed breastmilk, and the changes breastmilk undergoes during storage, remain unaddressed. Our research shows that the addition of food-derived compounds to breastmilk prior to freezing can delay the rancidification process. Moreover, compounds with multiple modes of action can be combined to inhibit fat breakdown and increase antioxidant capacity in breastmilk, maintaining milk quality during frozen storage for over 6 months. This technology has the potential to promote continued breastfeeding by addressing the unmet need for retaining breastmilk’s nutritional quality during storage. A prevailing theory concerning at-home breastmilk storage attributes rancidity to genetic variability in lipase levels.^55^ While we did not specifically test the lipase variation hypothesis in this study, our findings suggest that factors beyond lipase levels, such as MFG damage, phase separation, and antioxidant capacity, affect breastmilk quality during storage.

As noted by the WHO, breastfeeding is not simply a personal choice but rather a matter of public health and economic importance. The implications of our technology extend beyond improving milk storage as we aim to address long-standing economic and societal inequities negatively influencing parents, particularly women. By enabling parents to continue providing breastmilk to their infants, our technology supports the transition to a more sustainable food system, starting with infants’ first foods. Furthermore, by developing innovative, safe, and cost-effective solutions to breastmilk storage problems, our technology can help support lactating people’s participation in the workforce, which is critical for their economic empowerment and overall well-being.

## Acknowledgments

The authors thank Katie Silpe, Julie Valastyan, and the New Jersey Breastfeeding Coalition for general support of this work and soliciting milk donations; Shantal Garcia for assistance with the initial HTS screening protocol; and Itay Budin, Diane Stassi, and the PumpKin Baby Inc. team for helpful discussions during the writing process.

## Funding

The authors acknowledge financial support from the Jane Coffin Childs Memorial Fund for Medical Research and the Activate Fellowship (J.E.S.), Howard Hughes Medical Institute (J.E.S. and B.L.B.), and National Institutes of Health grant R41HD115506, USDA 2024-70436-42372 (J.E.S. and B.L.B.). The funders had no role in study design, data collection and analysis, decision to publish, or preparation of the manuscript.

## Competing interests

The authors disclose affiliation with and equity in PumpKin Baby Inc., a Princeton University spinout and for-profit public benefit corporation formed over the course of this work. PumpKin Baby Inc. is working to develop and commercialize the technology presented in this report, as the organization’s stated purpose is “to provide access to scientific research and products that aim to improve access to breastmilk, breastfeeding, and maternal and infant health.” Several patents related to the technology described in this manuscript are pending and assigned to Princeton University.

## Author Contributions (CRediT format)

**J.E.S.**: Conceptualization, Methodology, Investigation, Formal analysis, Data curation, Validation, Software, Writing - original draft, review, and editing, Project administration, Funding acquisition. Led the high-throughput screening campaign, developed and validated analytical assays, performed compound validation studies, and conducted biochemical analyses. **K.D.M.**: Formal analysis, Methodology, Data curation, Validation, Writing - review and editing, Performed statistical analyses of survey data and data interpretation. **B.L.B.**: Conceptualization, Investigation, Project administration, Supervision, Resources, Writing - original draft, review, and editing, Funding acquisition. Provided project oversight and strategic direction.

## Methods

### Study Design

This study was conducted in two phases, each independently approved by the Institutional Review Board (IRB) at Princeton University (IRB IDs: 16889 and 15531). Informed consent was obtained from all participants before their inclusion in the study. Phase 1 was focused on a cross-sectional nationwide survey to identify challenges associated with breastmilk storage practices. Phase 2 involved freshly expressed breastmilk collection to evaluate the influence of at-home storage practices on the nutritional quality of human milk.

### Phase 1: Survey of Breastmilk Storage Practices

The survey was conducted on a research platform generated and managed by Centiment Co (https://www.centiment.co/), which handled the deployment and data collection. Initially, 2,013 participants were screened and 1,049 were selected based on specified eligibility criteria identified within Centiment’s database. The questionnaire was available on the Centiment platform from May 30 to June 6, 2024, and included a series of yes/no and multiple-choice questions regarding demographics, breastmilk storage methods and durations, reasons for freezing breastmilk, and observed changes in milk quality post-storage. Additional questions addressed disposal of stored milk and challenges encountered. Original survey questions and exclusion criteria are accessible through Zenodo (refer to Data Availability).

### Phase 2: Collection of Human Milk Samples

Freshly-expressed human milk samples were collected from 14 lactating human donors. Participants were eligible to donate one fluid ounce (29.6 mL) of freshly pumped breastmilk if they met the following criteria: 1) Healthy breastfeeding mothers who were 18 to 45 years old; 2) Mothers who delivered a term infant (born 37 weeks or after); and 3) Mothers who exclusively breastfed for 12 or more weeks. Donors were recruited through advertisements in local parenting groups, social media, and lactation consultants. Each participant provided informed consent and was instructed on proper milk expression, collection, and storage techniques to ensure sample integrity. Samples were collected by the study personnel in the donor’s own storage container and transported to the laboratory on ice for immediate processing. After providing the breastmilk sample, participants received a USD 10 electronic gift card as compensation for their time and effort.

### Experimental Conditions and Storage Variables Analytical procedures

#### Microplate readers

BioTek Synergy Neo2 Multi-Mode readers with BioStack and BioSpa functionalities were used for lipase, lysozyme, protease, glycerol, FFA, and total antioxidant capacity measurements, as described below. Unless otherwise noted, fresh human milk served as the comparison standard, and water/DMSO was used as the vehicle control.

#### HTS screening

A multi-step HTS approach was developed to identify compounds that reduce lipolysis in human breastmilk during extended freezer storage. Briefly, 150 μL aliquots of freshly expressed (same-day) human milk, collected from a single donor were dispensed into 30 x 96-well plates (Corning 3903). Food and nutrient specific chemical libraries (MedChemExpress Food Additive Library, Food Sourced Compound Library, and Biolog Phenotype Microarrays PM1-5) were transferred into milk-containing wells using Scinomix 96 well pin replicators. The volume transfer from the pins was empirically determined to be ∼1.5 μL, leading to a 1% v/v treatment. Controls of 1% v/v DMSO and 1% v/v water were included in each plate. Plates were foil sealed and stored at -20 °C. At regular intervals for up to 1 week, plates were thawed and samples measured for lipolysis by transferring 2 μL of each treated milk sample into 96 well-plates containing a working solution (98% v/v, final concentration) of EnzChek Lipase Substrate (Thermo E33955). The working solution was prepared by dissolving 100 μg of substrate in 20 μL DMSO, followed by 5,000-fold dilution into PBS immediately prior to the experiment. Plates were incubated at 37 °C in the microplate reader for 2-6 hours, and Relative Fluorescence Units (λ_ex_ / λ_em_ of 500 nm / 530 nm) were measured. The method was validated using lipoprotein lipase from bovine milk (at 25 μg/mL with ≥2,000 units/mg, Sigma, L2254). Hit criteria were derived from Z-scores and defined as wells with RFUs ≤ 400.

#### Lysozyme

Lysozyme activity was assessed using the EnzCheck Lysozyme Assay Kit (Thermo E22013). A 1 mg/mL stock suspension of DQ lysozyme substrate was prepared by adding 1 mL deionized water to the lyophilized substrate. The working solution was prepared by diluting the stock suspension to 50 µg/mL in 1X reaction buffer (0.1 M sodium phosphate, 0.1 M NaCl, pH 7.5). Human milk samples (2 µL) were diluted with 48 µL reaction buffer in black 96-well plates (Corning 3904) and mixed with 50 µL of the working solution. Plates were incubated at 37 °C for 30 minutes protected from light. Fluorescence intensity was measured at λ_ex_ / λ_em_ of 485 nm / 530 nm using a microplate reader.

#### Protease

Protease activity was determined using the EnzCheck Protease Assay Kit (Thermo E6638). A 1 mg/mL stock solution of BODIPY FL casein substrate was prepared by adding 0.2 mL PBS to the lyophilized substrate. The working solution was prepared by diluting the stock solution to 10 µg/mL in 1X digestion buffer. Human milk samples (2 µL) were diluted with 48 µL digestion buffer in black 96-well plates (Corning 3904) and mixed with 50 µL of the working solution. Plates were incubated at 37 °C for 30 minutes protected from light. Proteolytic cleavage releases fluorescent BODIPY FL-labeled peptides, which were measured at λ_ex_ / λ_em_ of 500 nm / 535 nm using a microplate reader.

#### Glycerol

Glycerol content was quantified using the Glycerol-Glo Assay Kit (Promega J3151). Human milk samples (2 µL) were diluted with 48 µL glycerol lysis solution from the kit in white-walled 96-well plates (Corning 3903) and incubated for 30 minutes at 37 °C. The glycerol detection reagent was prepared by adding 10 µL reductase substrate per mL of glycerol detection solution and equilibrating for 60 minutes at room temperature. Kinetic enhancer (10 µL per mL) was then added to the detection reagent. An equal volume (50 µL) of this prepared glycerol detection reagent was added to each well containing the diluted milk samples. Plates were shaken for 30 seconds and incubated at room temperature for 60 minutes. Luminescence was measured using a microplate reader.

#### Total Antioxidant Capacity

Total antioxidant capacity of human milk was assessed using the Total Antioxidant Capacity Assay Kit with protein mask functionality (Abcam ab65329). The Cu^2+^ working solution was prepared by diluting Cu^2+^ Reagent 50x with Assay Buffer XXIV. Human milk samples (10 µL) were first diluted 1:1 with the protein mask reagent and then brought to a final volume of 100 µL, before adding 100 µL of the Cu^2+^ working solution. The plates were sealed and incubated at room temperature for 90 minutes on an orbital shaker protected from light prior to measuring absorbance at 570 nm using a microplate reader.

### Quantitation and statistical analyses

Software used to acquire and analyze data generated in this study consisted of: BioTek Gen5 for assessment of lipase, lysozyme, protease, glycerol, FFA, and total antioxidant capacity. GraphPad Prism10 was used for data visualization. Independent replicates are defined as milk samples obtained from the same donor that were individually prepared, stored in separate tubes, measured in distinct wells, and processed on the same day. Unless otherwise noted, data are presented as the means ± std. Where indicated, biological replicates (Fig 2c) are defined as milk samples obtained from different donors, and analyses carried out on the same day.

### Data availability statement

Data reported in this study are provided as a Source Data file. All materials associated with this study have been deposited on Zenodo (doi: 10.5281/zenodo.14252924). Other experimental data that support the findings of this study are available without restriction by request from the corresponding author.

---

**Supplemental Figure 1:**
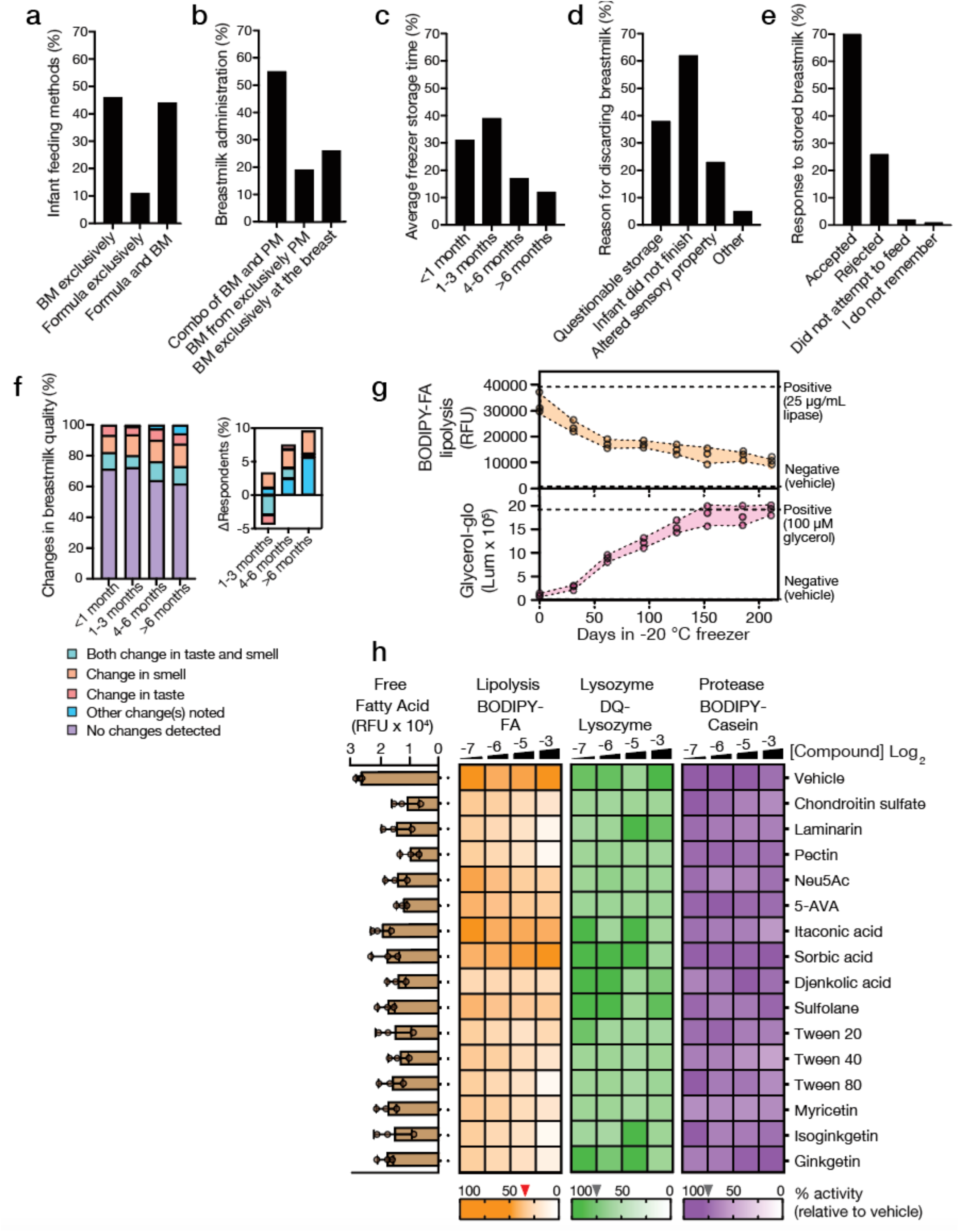
Survey Data and Biochemical Assays Related to Breastmilk Storage and Preservation. **(a-f):** Survey results from 1,049 respondents on milk storage practices. **(a)** Infant feeding methods in first 6 months: exclusive breastmilk (*n* = 479), exclusive formula (*n* = 111), or combined (*n* = 459). **(b)** Breastmilk administration: combined breastfeeding and pumped milk (*n* = 514), exclusive breastfeeding (*n* = 241), or pumped milk only (*n* = 178). BM: breastfed breastmilk; PM: pumped breastmilk. **(c)** Freezer storage duration: <1 month (*n* = 224), 1-3 months (*n* = 283), 4-6 months (*n* = 122), >6 months (*n* = 89). **(d)** Milk discard reasons: storage concerns (*n* = 293), incomplete feeding (*n* = 486), sensory changes (*n* = 180). **(e)** Infant acceptance of frozen milk: accepted (*n* = 701), rejected (*n* = 269, inclusive of occasional and outright rejection), not attempted (*n* = 17), no recall (*n* = 15). **(f)** Quality changes over storage time, with the inset showing the percent change from the earlier time point. **(g)** Storage-dependent changes at -20 °C: BODIPY-FA lipolysis activity (upper) and glycerol accumulation (lower). Dashed lines: controls (positive: 25 μg/mL lipase or 100 μM glycerol; negative: vehicle). **(h)** Validation of HTS hits at a single concentration (Log_2_(-3)) on FFA levels and, in a concentration-dependent manner, on lipolysis, lysozyme, and protease activities. Compound concentrations and abbreviations as in Figure 1b. Red triangle: normalized lipolysis hit threshold; gray triangles: normalized lysozyme and protease counter thresholds, each relative to vehicle-treated milk. Data are shown as means ± std (g, h-bars) or median of triplicates (h-heatmaps).

**Supplemental Table 1:** Survey Demographics.

|  |  |
| --- | --- |
| **Age** | **Mean ± SD** |
|  | 30.6 ± 6.7 |
| **Race** | **% (n)** |
| White | 61% (644) |
| Black or African American | 23% (239) |
| Hispanic or Latino | 17% (176) |
| Asian | 4% (44) |
| American Indian or Alaska Native | 2% (24) |
| Native Hawaiian or Pacific Islander | 1% (12) |
| Middle Eastern or North African (MENA) | 1% (7) |
| Prefer not to answer | <1% (3) |
| Other | 1% (10) |
| **Employment Status** | **% (n)** |
| Full-time employed | 44% (438) |
| Unemployed | 22% (222) |
| Part-time employed | 19% (187) |
| Student (independent of other employment) | 4% (43) |
| Prefer not to answer | 2% (23) |
| Retired | 0.3% (3) |
| Other | 8% (78) |
| **Household Income** | **% (n)** |
| \$0-\$19,999 | 14% (n) |
| \$20,000-\$49,999 | 34% (n) |
| \$50,000-\$99,999 | 31% (324) |
| \$100,000-\$199,999 | 16% (164) |
| \$200,000+ | 4% (47) |
| Not sure | 0% (5) |
| Prefer not to answer | 1% (8) |
| **Infant Birth Year** | **% (n)** |
| 2020 | 16.9% (177) |
| 2021 | 18% (184) |
| 2022 | 25% (158) |
| 2023 | 27% (284) |
| 2024 | 14% (146) |

**Supplemental Table 2:** Breastmilk Storage Practices Among Participants.

|  |  |
| --- | --- |
| **Ever stored expressed breastmilk** | **% (n)** |
| Yes | 100% (1049) |
| No | 0% |
| **Amount of expressed breastmilk stored** | **% (n)** |
| Some | 39.0% (410) |
| About a half | 41% (425) |
| Most | 13% (134) |
| All | 6% (63) |
| I don't remember | 2% (17) |
| **Storage methods of expressed breastmilk** | **% (n)** |
| Room temperature | 10% (108) |
| Refrigerator | 64% (673) |
| Freezer | 68% (718) |
| Unsure | 0.2% (3) |
| Other | 0.1% (2) |
| **Reason for freezing expressed breastmilk** | **% (n)** |
| Work | 38% (271) |
| Travel | 23% (162) |
| Oversupply | 63% (451) |
| Daycare | 24% (175) |
| Other | 9% (66) |
| **Average storage time of breastmilk in the freezer** | **% (n)** |
| Less than 1 month | 31.2% (224) |
| 1-3 months | 39% (283) |
| 4-6 months | 17% (122) |
| More than 6 months | 12% (89) |
| **Changes in breastmilk quality after defrosting** | **% (n)** |
| Change in smell | 13% (93) |
| Change in taste | 6% (45) |
| Change in taste and smell | 10% (71) |
| No changes detected | 69% (498) |
| Other | 2% (11) |
| **Average storage time of breastmilk in the refrigerator** | **% (n)** |
| Less than 24 hours | 56% (168) |
| 1 to 3 days | 40% (121) |
| 3 to 5 days | 3% (9) |
| More than 5 days | 1% (3) |
| **Changes in breastmilk quality after refrigerator storage** | **% (n)** |
| Change in smell | 14% (41) |
| Change in taste | 7% (21) |
| Change in taste and smell | 10% (30) |
| No changes detected | 67% (203) |
| Other | 2% (6) |
| **Infant response to stored breastmilk** | **% (n)** |
| Accepted | 70% (701) |
| Sometimes rejected | 23% (235) |
| Rejected | 3% (34) |
| Did not attempt to feed | 2% (17) |
| I do not remember | 1% (15) |
| **Reasons for discarding breastmilk** | **% (n)** |
| Questionable storage method (e.g. power outage) | 38% (293) |
| Infant did not finish the feeding | 62% (486) |
| The milk smelled off | 23% (180) |
| Other | 5% (40) |

## Notes

### Summary of Updates

There was a formatting issue in the introduction that previously read "8's work-life system" but should read "In today's work-life system".

## References

1. Nardella, D., Canavan, M., Sharifi, M., and Taylor, S. (2024). Quantifying the Association between Pump Use and Breastfeeding Duration. J. Pediatr. 274, 114192. 10.1016/j.jpeds.2024.114192.

2. CDC (2023). 2022 Breastfeeding Report Card. Cent. Dis. Control Prev. https://www.cdc.gov/breastfeeding/data/reportcard.htm.

3. CDC (2024). Facts About Nationwide Breastfeeding Goals. Cent. Dis. Control Prev. https://www.cdc.gov/breastfeeding/data/facts.html.

4. Gao, G., and Livingston, G. Working while pregnant is much more common than it used to be. Pew Res. Cent. https://www.pewresearch.org/short-reads/2015/03/31/working-while-pregnant-is-much-more-common-than-it-used-to-be/.

5. Light, P. (2013). Why 43% of Women With Children Leave Their Jobs, and How to Get Them Back. The Atlantic. https://www.theatlantic.com/sexes/archive/2013/04/why-43-of-women-with-children-leave-their-jobs-and-how-to-get-them-back/275134/.

6. Women’s salaries plummet after giving birth. Here’s one way to restore their earning power (2020). USC Today. https://news.usc.edu/women-giving-birth-child-penalty-salary-gap-usc-research/.

7. Nations, U. Breastfeeding and Work: A Balancing Act. U. N. https://www.un.org/en/un-chronicle/breastfeeding-and-work-balancing-act.

8. Chen, Y.C., Wu, Y.-C., and Chie, W.-C. (2006). Effects of work-related factors on the breastfeeding behavior of working mothers in a Taiwanese semiconductor manufacturer: a cross-sectional survey. BMC Public Health 6, 160. 10.1186/1471-2458-6-160.

9. Webb, B.H., and Hall, S.A. (1935). Some Physical Effects of Freezing upon Milk and Cream. J. Dairy Sci. 18, 275–286. 10.3168/jds.S0022-0302(35)93146-0.

10. Ballard, O., and Morrow, A.L. (2013). Human Milk Composition: Nutrients and Bioactive Factors. Pediatr. Clin. North Am. 60, 49. 10.1016/j.pcl.2012.10.002.

11. Liang, N., Koh, J., Kim, B.J., Ozturk, G., Barile, D., and Dallas, D.C. (2022). Structural and functional changes of bioactive proteins in donor human milk treated by vat-pasteurization, retort sterilization, ultra-high-temperature sterilization, freeze-thawing and homogenization. Front. Nutr. 9, 926814. 10.3389/fnut.2022.926814.

12. García-Lara, N.R., Escuder-Vieco, D., García-Algar, O., De la Cruz, J., Lora, D., and Pallás-Alonso, C. (2012). Effect of Freezing Time on Macronutrients and Energy Content of Breastmilk. Breastfeed. Med. 7, 295–301. 10.1089/bfm.2011.0079.

13. Hamosh, M., Ellis, L.A., Pollock, D.R., Henderson, T.R., and Hamosh, P. (1996). Breastfeeding and the Working Mother: Effect of Time and Temperature of Short-term Storage on Proteolysis, Lipolysis, and Bacterial Growth in Milk. Pediatrics 97, 492–498. 10.1542/peds.97.4.492.

14. Ahrabi, A.F., Handa, D., Codipilly, C.N., Shah, S., Williams, J.E., McGuire, M.A., Potak, D., Aharon, G.G., and Schanler, R.J. (2016). Effects of Extended Freezer Storage on the Integrity of Human Milk. J. Pediatr. 177, 140–143. 10.1016/j.jpeds.2016.06.024.

15. Hung, H.-Y., Hsu, Y.-Y., Su, P.-F., and Chang, Y.-J. (2018). Variations in the rancid-flavor compounds of human breastmilk under general frozen-storage conditions. BMC Pediatr. 18, 94. 10.1186/s12887-018-1075-1.

16. Braune, K., Kraemer, L.A., Weinstein, J., Zayani, A., and Heinemann, L. (2019). Storage Conditions of Insulin in Domestic Refrigerators and When Carried by Patients: Often Outside Recommended Temperature Range. Diabetes Technol. Ther. 21, 238–244. 10.1089/dia.2019.0046.

17. Francis, J., and Dickton, D. (2018). Feeding and refusal of expressed and stored human (FRESH) milk study - a short communication. Health Kinesiol. Fac. Publ. Present.

18. Hsiao-Ying Hung, Y. Hsu, P. Su, and Ying-Ju Chang (2018). Variations in the rancid-flavor compounds of human breastmilk under ge neral frozen-storage conditions. BMC Pediatr.

19. T. Olivecrona and O. Hernell (1976). Human milk lipases and their possible role in fat digestion. Pädiatr. Pädologie.

20. J. Spitzer, K. Klos, and A. Buettner (2013). Monitoring aroma changes during human milk storage at +4 °C by sensory and quantification experiments. Clin. Nutr.

21. S. Duncan, G. L. Christen, and M. P. Penfield (1991). Rancid flavor of milk: relationship of acid degree value, free fatty a cids, and sensory perception.

22. Eglash, A., Simon, L., and Academy of Breastfeeding Medicine (2017). ABM Clinical Protocol #8: Human Milk Storage Information for Home Use for Full-Term Infants, Revised 2017. Breastfeed. Med. Off. J. Acad. Breastfeed. Med. 12, 390–395. 10.1089/bfm.2017.29047.aje.

23. Bank, M.R., Kirksey, A., West, K., and Giacoia, G. (1985). Effect of storage time and temperature on folacin and vitamin C levels in term and preterm human milk. Am. J. Clin. Nutr. 41, 235–242. 10.1093/ajcn/41.2.235.

24. Romeu-Nadal, M., Castellote, A.I., and López-Sabater, M.C. (2008). Effect of cold storage on vitamins C and E and fatty acids in human milk. Food Chem. 106, 65–70. 10.1016/j.foodchem.2007.05.046.

25. Buss, I., McGill, F., and Winterbourn, C. Vitamin C is reduced in human milk after storage.

26. Winters, M., and Tompkins, C.A. (1936). A Pectin-Agar Preparation for Treatment of Diarrhea of Infants. Am. J. Dis. Child. 52, 259–265. 10.1001/archpedi.1936.04140020002001.

27. EFSA Panel on Food Additives and Flavourings (FAF), Younes, M., Aquilina, G., Castle, L., Engel, K.-H., Fowler, P., Frutos Fernandez, M.J., Fürst, P., Gürtler, R., Husøy, T., et al. (2021). Opinion on the re-evaluation of pectin (E 440i) and amidated pectin (E 440ii) as food additives in foods for infants below 16 weeks of age and follow-up of their re-evaluation as food additives for uses in foods for all population groups. EFSA J. 19, e06387. 10.2903/j.efsa.2021.6387.

28. Donadio, J.L.S., Fabi, J.P., Sztein, M.B., and Salerno-Gonçalves, R. (2024). Dietary fiber pectin: challenges and potential anti-inflammatory benefits for preterms and newborns. Front. Nutr. 10, 1286138. 10.3389/fnut.2023.1286138.

29. García-Lara, N.R., Vieco, D.E., De la Cruz-Bértolo, J., Lora-Pablos, D., Velasco, N.U., and Pallás-Alonso, C.R. (2013). Effect of Holder Pasteurization and Frozen Storage on Macronutrients and Energy Content of Breast Milk. J. Pediatr. Gastroenterol. Nutr. 57, 377. 10.1097/MPG.0b013e31829d4f82.

30. Ahrabi, A.F., Handa, D., Codipilly, C.N., Shah, S., Williams, J.E., McGuire, M.A., Potak, D., Aharon, G.G., and Schanler, R.J. (2016). Effects of Extended Freezer Storage on the Integrity of Human Milk. J. Pediatr. 177, 140–143. 10.1016/j.jpeds.2016.06.024.

31. H. Lev, A. Ovental, D. Mandel, F. Mimouni, R. Marom, and R. Lubetzky (2014). Major losses of fat, carbohydrates and energy content of preterm human milk frozen at −80°C. J. Perinatol.

32. Kaya, Ö., and Çınar, N. (2023). The effects of freezing and thawing on mature human milk’s contains: A systematic review. Midwifery 118, 103519. 10.1016/j.midw.2022.103519.

33. Orbach, R., Mandel, D., Mangel, L., Marom, R., and Lubetzky, R. (2019). The Effect of Deep Freezing on Human Milk Macronutrients Content. Breastfeed. Med. 14, 172–176. 10.1089/bfm.2018.0226.

34. Schlotterer, H.R., and Perrin, M.T. (2018). Effects of Refrigerated and Frozen Storage on Holder-Pasteurized Donor Human Milk: A Systematic Review. Breastfeed. Med. Off. J. Acad. Breastfeed. Med. 13, 465–472. 10.1089/bfm.2018.0135.

35. Lina Zhang, Yanyan Wu, Yaping Ma, Zhuangjian Xu, Ying Ma, and P. Zhou (2020). Macronutrients, total aerobic bacteria counts and serum proteome of hu man milk during refrigerated storage.

36. Yu-Chuan Chang, Chao-Huei Chen, and Ming-Chih Lin (2012). The macronutrients in human milk change after storage in various conta iners. Pediatr. Neonatol.

37. Lugonja, N., Marinković, V., Miličić, B., Avdalović, J., Vrvić, M., and Spasić, S. (2023). Effect of storage process on nutritive properties of preterm human milk. Chem. Ind. Chem. Eng. Q. 29, 141–148.

38. Lopes, L.M.P., Chaves, J.O., Cunha, L.R.D., Passos, M.C., and Menezes, C.C. (2020). Hygienic-sanitary quality and effect of freezing time and temperature on total antioxidant capacity of human milk. Braz. J. Food Technol. 23, e2019179. 10.1590/1981-6723.17919.

39. Ribeiro, V.P.D., Tinoco, R.B., Chamon, A.L.B., Pessoa, I.S., Santos, T.C.D., Silva, R.S., and Fronza, M. (2023). The Influence of Time and Temperature on Human Milk Storage Antioxidant Properties, Oxidative Stress, and Total Protein. J. Hum. Lact. Off. J. Int. Lact. Consult. Assoc. 39, 308–314. 10.1177/08903344221126669.

40. Sheen, W., Ahmed, M., Patel, H., Codipilly, C.N., and Schanler, R.J. (2021). Is the Antioxidant Capacity of Stored Human Milk Preserved? Breastfeed. Med. Off. J. Acad. Breastfeed. Med. 16, 564–567. 10.1089/bfm.2020.0407.

41. Li, N., Huang, G., Zhang, Y., Zheng, N., Zhao, S., and Wang, J. (2022). Diversity of Volatile Compounds in Raw Milk with Different n-6 to n-3 Fatty Acid Ratio. Anim. Open Access J. MDPI 12, 252. 10.3390/ani12030252.

42. Jiang, S., Luo, W., Peng, Q., Wu, Z., Li, H., Li, H., and Yu, J. (2022). Effects of Flash Evaporation Conditions on the Quality of UHT Milk by Changing the Dissolved Oxygen Content in Milk. Foods 11, 2371. 10.3390/foods11152371.

43. Ghosh, S., and Coupland, J.N. (2008). Factors affecting the freeze–thaw stability of emulsions. Food Hydrocoll. 22, 105–111. 10.1016/j.foodhyd.2007.04.013.

44. Degner, B.M., Chung, C., Schlegel, V., Hutkins, R., and McClements, D.J. (2014). Factors Influencing the Freeze-Thaw Stability of Emulsion-Based Foods. Compr. Rev. Food Sci. Food Saf. 13, 98–113. 10.1111/1541-4337.12050.

45. O’Flynnl, B.G., and Mittagl, T. (2021). The role of liquid-liquid phase separation in regulating enzyme activity. Curr. Opin. Cell Biol. 69, 70. 10.1016/j.ceb.2020.12.012.

46. Digvijay, Kelly, A.L., and Lamichhane, P. (2023). Ice crystallization and structural changes in cheese during freezing and frozen storage: implications for functional properties. Crit. Rev. Food Sci. Nutr., 1–24. 10.1080/10408398.2023.2277357.

47. Grebenteuch, S., Kroh, L.W., Drusch, S., and Rohn, S. (2021). Formation of Secondary and Tertiary Volatile Compounds Resulting from the Lipid Oxidation of Rapeseed Oil. Foods 10, 2417. 10.3390/foods10102417.

48. Cervantes-Paz, B., Ornelas-Paz, J.D.J., Ruiz-Cruz, S., Rios-Velasco, C., Ibarra-Junquera, V., Yahia, E.M., and Gardea-Béjar, A.A. (2017). Effects of pectin on lipid digestion and possible implications for carotenoid bioavailability during pre-absorptive stages: A review. Food Res. Int. 99, 917–927. 10.1016/j.foodres.2017.02.012.

49. Romeu-Nadal, M., Castellote, A.I., and López-Sabater, M.C. (2008). Effect of cold storage on vitamins C and E and fatty acids in human milk. Food Chem. 106, 65–70. 10.1016/j.foodchem.2007.05.046.

50. Pehrsson, P.R., Patterson, K.Y., and Khan, M.A. (2014). Selected vitamins, minerals and fatty acids in infant formulas in the United States. J. Food Compos. Anal. 36, 66–71. 10.1016/j.jfca.2014.06.004.

51. Heck, A.M., Yanovski, J.A., and Calis, K.A. (2000). Orlistat, a New Lipase Inhibitor for the Management of Obesity. Pharmacother. J. Hum. Pharmacol. Drug Ther. 20, 270–279. 10.1592/phco.20.4.270.34882.

52. Smith, J.P., Baker, P., Mathisen, R., Long, A., and Waring, M. A proposal to recognize investment in breastfeeding as a carbon offset. Bull. World Health Organ.

53. Smith, J.P. (2013). “Lost milk?”: Counting the economic value of breast milk in gross domestic product. J. Hum. Lact. Off. J. Int. Lact. Consult. Assoc. 29, 537–546. 10.1177/0890334413494827.

54. Nelson, A. The Politics of Breastfeeding (And Why It Must Change). Forbes. https://www.forbes.com/sites/amynelson1/2018/10/24/the-politics-of-breastfeeding-and-why-it-must-change/.

55. High Lipase Milk: Cause, Effects, and How to Manage (2020). Healthline. https://www.healthline.com/health/breastfeeding/high-lipase-milk.

